# Forced exercise modulates retinal inflammatory response and regulates miRNA expression to promote retinal neuroprotection during degeneration

**DOI:** 10.1101/2025.07.16.665187

**Authors:** Hayden Haupt, Vivian S. Chen, Teele Palumaa, Teresa E. Anderson, Gabriela Sanchez Rodriguez, Joshua Chu-Tan, Riccardo Natoli, Andrew J. Feola, John M. Nickerson, Machelle T. Pardue, Jeffrey H. Boatright, Katie L. Bales

**Affiliations:** Atlanta VA Center for Visual and Neurocognitive Rehabilitation, Decatur, GA; Department of Biomedical Engineering, Georgia Institute of Technology, Atlanta, GA; Department of Ophthalmology, Emory University, Atlanta, GA; Institute of Genomics, University of Tartu, Tartu, Estonia; School of Electrical and Computer Engineering, Georgia Institute of Technology, Atlanta GA; Eccles Institute of Neuroscience, John Curtin School of Medical Research, College of Health and Medicine, The Australian National University, Acton, Australia; School of Medicine and Psychology, College of Health and Medicine, The Australian National University, Acton, Australia

**Author notes:** Corresponding Author: Jeffrey Boatright, Decatur, GA 30033, USA. <u>Support:</u> VA BLR&D IK2BX005304, VA RR&D C9246C, VA RRD IK6 RX003134, NIH R21 EY027005, NIH P30EY06360, The Challenge Grant (Research to Prevent Blindness), The Abraham J. & Phyllis Katz Foundation, Atlanta Veterans Education and Research (FAVER) Foundation, European Union Horizon Europe research and innovation program under the Marie Skłodowska-Curie grant agreement No 101153901. <u>Commercial relationships:</u> none.

**Keywords:** Exercise, Retinal Neuroprotection, miRNA, Retinal inflammation

## Abstract

**Background:** Our labs have demonstrated exercise is protective in animal models of retinal degeneration (RD). Inflammation drives RD progression, and is regulated by the recruitment and reactivity of glia cells as well as through small non-coding RNAs, microRNAs (miRNAs). Here, we explore the effects of treadmill exercise on the recruitment and reactivity of retinal inflammatory cells within the neural retina and miRNA expression in a light-induced retinal degeneration model (LIRD) that exhibits phenotypes found in patients with RD.

**Methods:** Male 6-week-old BALB/c mice were randomly assigned to either active or inactive groups. Active groups were exercised by treadmill 1 hour a day for two weeks at a speed of 10m/min, meanwhile inactive groups were placed on static treadmills for the same duration. Light induced retinal degeneration (LIRD) was induced during the second week of exercise using light exposure of 5000 lux, control animals were kept at 50 lux. Retinal function was assessed using electroretinography (ERG) 5 days after LIRD. Retinas were collected 1-day and 5-days post-LIRD, sagittal sections were stained for inflammatory markers (GFAP and Iba1), TUNEL (cell death), and photoreceptor nuclei (outer nuclear layer; ONL) were quantified. RNA was extracted and miRNA expression quantified with GeneChip miRNA 4.0 array.

**Results:** Active+LIRD mice demonstrated significant preservation of retinal function, evidenced by higher a-wave and b-wave amplitudes in ERG 5-days post-LIRD, compared to inactive+LIRD mice. Retinal sections from active+LIRD mice had fewer Iba1+ cells and decreased GFAP labeling 5-days post-LIRD compared to inactive+LIRD mice. Active+LIRD mice had fewer ONL TUNEL+ cells compared to inactive+LIRD mice. Inactive+LIRD mice showed a decline in ONL counts 1-day post-LIRD with significant loss 5-days post-LIRD compared to active+LIRD mice. In active groups, exercise promoted significant differences in miRNA expression, such as miR-302b, miR-192-5p, miR-187 compared to inactive groups.

**Conclusions:** Our results indicate that treadmill exercise preserved photoreceptor density, slowed and or prevented apoptosis in the ONL, and decreased the presence/recruitment of inflammatory cells in the neural retina. Altered miRNA expression profiles in active groups are associated with cell survival (miR-302b), oxidative stress regulation (miR-192-5p) and photoreceptor homeostasis (miR-187). These results reveal how exercise alters the retinal inflammatory response over the course of 1-day to 5-days, providing insight into exercise-based therapies and treatments for RD and neuroinflammatory diseases.

## Introduction

Physical exercise has been shown to be an effective, non-invasive approach to halt and potentially prevent neurodegenerative disease progression. Our research group and others have investigated the neuroprotective effects of exercise in many animal models of retinal degenerative diseases^1–5^. We have shown exercise reduced retinal pigmented epithelium (RPE) stress, reduced photoreceptor cell death, preserved retinal structure and function, increased retinal astrocyte plasticity and reduced the expression of proinflammatory chemokines keratinocyte-derived chemokine (KC) and interferon gamma inducible protein-10 (IP-10)^3–6^.

One of the major targets for managing degenerative diseases is modulating the inflammatory response associated with disease onset and progression. Neuroinflammation is a hallmark feature of retinal degenerative diseases, such as age-related macular degeneration and retinitis pigmentosa^7,8^. Physical exercise has been shown to regulate fundamental inflammatory pathways through modulating pro-inflammatory cytokine profiles, which result in dampening of the innate immune response^2^. The regulation of inflammatory response is partly mediated by microRNAs (miRNAs), which modulate the expression of genes involved in cytokine production, immune cell activation, and resolution of inflammation^9^.

To investigate how exercise influences retinal inflammatory cell response, photoreceptor cell apoptosis, and miRNA expression we have incorporated the light-induced retinal degeneration model (LIRD). Through this model we are able to assess how retinal structure, function and molecular markers are altered in the initial onset of retinal degeneration by assessing 1-day post-LIRD as well as 5-days post-LIRD. LIRD is a powerful model that recapitulates several phenotypes associated with retinal degenerative diseases and allows the precise control of retinal degeneration onset and intensity^10^. By incorporating this inducible model as well as examining 1- and 5-days post LIRD, we are able to directly measure changes in glial inflammatory response, layer-specific photoreceptor survival, and expression of regulatory miRNAs in mice retinas in response to forced treadmill exercise at specific neurodegeneration checkpoints.

## Methods

### Animals

All animal procedures were approved by the Atlanta VA Institutional Animal Care and Use Committee and conform to the ARVO Statement for the Use of Animals in Ophthalmic and Vision Research. Adult BALB/c male mice were purchased from Charles River (8–10 weeks old; Wilmington, MA, USA) and housed under a 12:12 light: dark cycle. During the light cycle, light levels measured at the floor of the mouse cages ranged from 5 to 45 lux. Mice had access to standard mouse chow (Teklad Global 18% Protein Rodent Diet 2918, Irradiated, Rockville, MD) and water ad libitum.

### Experimental design

Mice were randomly assigned to one of the following four groups: inactive + dim (*n* = 24), active + dim (*n* = 24), inactive + LIRD (*n* = 24), and active + LIRD (*n* = 24). Active groups ran on a rodent treadmill once daily at 10 m per minute (m/min), 5 days per week for 2 weeks. Inactive groups were placed on a static treadmill for the same amount of time. On the day of LIRD, mice were exposed to toxic light within 30 min of the end of the treadmill session. Mice included in the 1-day post-LIRD timepoint were euthanized following their exercise regimen 24 hours following light damage. Mice utilized for the 5-day post-LIRD timepoint were given an additional week of treadmill running, electroretinography (ERG) was performed to assess retinal function. Following 2 final days of exercise, mice were exercised in a staggered fashion so that each mouse could be euthanized immediately after the end of the 1 h treadmill session. Mice were euthanized via CO_2_ gas inhalation and secondary cervical dislocation, eyes were enucleated for retinal flat mounts and retinal astrocyte isolation.

### Exercise regimen and light exposure

In accordance with previous studies, active mice ran 60 min per day between ZT3-5 on treadmills equipped with electric shock gratings (Exer-3/6; Columbus Instruments, Columbus, OH, USA). Grating inactivated if the mice received 10 shocks in a single session. Before daily treadmill sessions began, the mice were trained for 5 to 10 min at a pace of 5 to 10 m/min for 2 consecutive days on the treadmill. The mice learned to maintain running within the first few days of treadmill exposure, receiving very few shocks after the first 1 to 2 days. Inactive mice were placed on static treadmills. Following 2 weeks of exercise, the mice were exposed to typical laboratory lighting (50 lux; dim) or toxic light (5000 lux; LIRD) for 4 h using a white light emitting diode (LED) light panel (LED500A; Fancierstudio, Hayward, CA, USA). This level of toxic light is a moderate brightness to induce retinal degeneration. For light exposure, animals were individually housed in shoebox containers with the LED light panel placed above as previously described^2^. Room and light box temperatures were closely monitored to ensure animal welfare. Following light damage, mice were exercised for an additional week and were euthanized 1-day and 5-days post-light exposure.

### Electroretinography (ERG)

Retinal function was measured with a commercial electroretinography (ERG) system (Bigshot; LKC Technologies, Gaithersburg, MD). After overnight dark adaptation, mice were anesthetized (ketamine [80 mg/kg]/xylazine [16 mg/kg]). All procedures were performed under dim red light during the subjective day. The pupils were dilated (1% tropicamide; Alcon Laboratories, Ft. Worth, TX) and corneas were anesthetized (1% tetracaine; Alcon Laboratories, Ft. Worth, TX). Body temperature was maintained with a heating pad at 37°C (ATC 1000; World Precision Instruments, Sarasota, FL) for the duration of the session. The ERG protocol consisted of a ten-step series of full-field flash stimuli produced by a Ganzfeld dome under both dark-adapted scotopic conditions (-3.0 to 2.1 log cd s/m^2^) to test rod and rod/cone pathways and light-adapted photopic conditions to test cone pathway function (2.0 log cd s/m^2^ presented as 6.1 Hz flicker with a 30 cd/m^2^ background light). Custom gold-loop wire electrodes were placed on the center of each eye through a layer of 1% methylcellulose to measure the electrical response of the eye to each flash. Reference and ground platinum needle reference electrodes (1cm; Natus Medical Incorporated, Pleasanton, CA) were inserted subcutaneously in cheeks and the tail, respectively. Once ERG was completed, mice were given an intraperitoneal (IP) injection of atipamezole (1 mg/kg; Antisedan, Zoetis, Parsippany, NJ) to counteract the effects of xylazine, administered saline eye drops and allowed to recover on a heating pad (37°C) before being returned to housing. ERG responses from both eyes were averaged.

### Histology

Eyes enucleated for histological analysis and were fixed in 97% methanol/ 3% acetic acid, dehydrated, embedded in paraffin, and sectioned through the sagittal plane on a microtome (5μm). Retinal sections were cut superior to inferior sections (0.5 μm) of the retina bisecting the optic disc using a rotary microtome in the superior–inferior plane, bisecting the optic nerve. Retinal spidergrams were constructed by plotting the number of photoreceptor nuclei present in the outer nuclear layer (ONL) as a function of position in the retina relative to the optic nerve. ONL nuclei were counted in a semiautomated fashion using Adobe Photoshop (24.0.0 release) to positively identify and count them within 100-μm-wide segments spaced at 250, 750, and 1,250μm from the optic nerve head in the superior and inferior directions.

### Immunofluorescence

Eyes were fixed in 97% methanol/ 3% acetic acid, dehydrated, embedded in paraffin, and sectioned through the sagittal plane on a microtome (5μm). Sections containing the optic nerve were selected for staining to ensure that consistent regions were examined between animals. The slides were deparaffinized across five Coplin jars with 100 mL of xylene for 2 min each, consecutively. Then the slides were rehydrated in a series of 100 mL ethanol solutions for 2 min each: 100%, 90%, 80%, 70%, 60%, and 50%. Following rehydration, retinal sections were blocked for 30 minutes at room temperature in 5% normal donkey serum in PBS with 0.01% sodium azide and 0.3% Triton X-100. Primary antibodies were diluted in 5% normal donkey serum in PBS with 0.01% sodium azide and incubations were performed overnight at 4°C using GFAP antibody (ab53554, abcam, Cambridge, United Kingdom, 1:100 dilution) and Iba1 antibody (ab178847, abcam, Cambridge, United Kingdom, 1:100 dilution). Secondary antibodies were diluted in PBS and incubations were performed for 1 hour at room temperature in the dark (Alexa Fluor ® 568 donkey anti-goat; 1:500, A11057, ThermoFisher Scientific, Waltham, MA; Alexa Fluor ® 647 donkey anti-rabbit; 1:500, A31573, ThermoFisher Scientific, Waltham, MA). Retinal sections were mounted with ProLong ® Gold Antifade Reagent with DAPI (#8961, Cell Signaling Technology Inc., Danvers, MA). For immunofluorescence negative controls, retinal flat mounts from each experimental group were incubated with primary antibody diluent alone with no primary antibody, followed by incubation with secondary antibody to ensure staining was produced from the detection of the antigen by the primary antibody. Retinal sections were imaged using a Nikon A1R HD25 confocal microscope with a Plano Apo 20x NA 0.75 objective and compiled and quantified using ImageJ software. GFAP and Iba1 labeling quantification are the result of positive GFAP and Iba1 labeling throughout the retinal layers.

### MicroArray

For miRNA expression analyses, retinas were collected in the middle of the light phase (1 day after LIRD: inactive+dim, n=5; active+dim, n=5; inactive+LIRD, n=; active+LIRD, n=5); 5 days after LIRD: inactive+dim, n=5; active+dim, n=4; inactive+LIRD, n=; active+LIRD, n=5, and placed in stabilization reagent (TRIzol, Ambion, Carlsbad, California; Catalog no. 15596018). Total RNA was extracted (RNeasy, QIAGEN; Catalog no. 74106) following the manufacturer’s protocol. RNA quality was analyzed with Bioanalyzer (Agilent, Santa Clara, CA) and samples with RIN values >7 were used. Samples were submitted to Tempus Multi-Omics for further processing and analysis. miRNA expression was analyzed using the Affymetrix GeneChip miRNA 4.1 Array. Raw CEL files were imported into Transcriptome Analysis Console (TAC) 4.0 (Thermo Fisher Scientific). Data was normalized to account for chip-related batch effects. Robust multiarray average (RMA) was used for background correction, normalization and miRNA expression calculation. Differential expression analysis was conducted separately for retinas collected 1-day and 5-days post-LIRD. Comparisons were made between the exercise groups (exercised vs. control), light exposure conditions (dim vs. LIRD), and their interaction. The results were filtered for mouse miRNAs. A *p* value of <0.05 was considered statistically significant.

### Masking and Statistical analysis

Sample size was determined based on our previously reported data^3,4,11^. Researchers who analyzed the data were blinded to the experimental procedures and were masked to the specific treatment groups from which sampling arose. All data are presented as mean ± standard error of the mean (SEM). Statistical analyses were performed using Graphpad Prism 10.2.3 (San Diego, CA, USA). Two-way ANOVAs dividing groups by exercise and light exposure to test the interactive effects of the independent variables and Tukey’s multiple comparison tests were performed. All *p*-values lower than .05 were considered statistically significant. The ROUT method (with Q set to 1%) was used to detect outliers.

## Results

### Active+LIRD mice have conserved retinal function 5-days post-LIRD

Electroretinograms were performed on all experimental groups to assess retinal function in order to confirm treadmill exercise preserved retinal function (**Figure 1A, B**). As we have previously shown, active+LIRD mice undergoing retinal degeneration revealed statistically significant preservation of retinal function 5-days post-LIRD with 1.98x (p<0.0093) greater scotopic a-wave amplitudes and 1.72x (p<0.0004) greater b-wave amplitudes compared to inactive+LIRD mice (two-way ANOVA with Tukey’s multiple comparison analysis was performed, n=10 per group, average of both eyes). These data support and confirm that treadmill exercise preserved retinal function.

**Figure 1.**
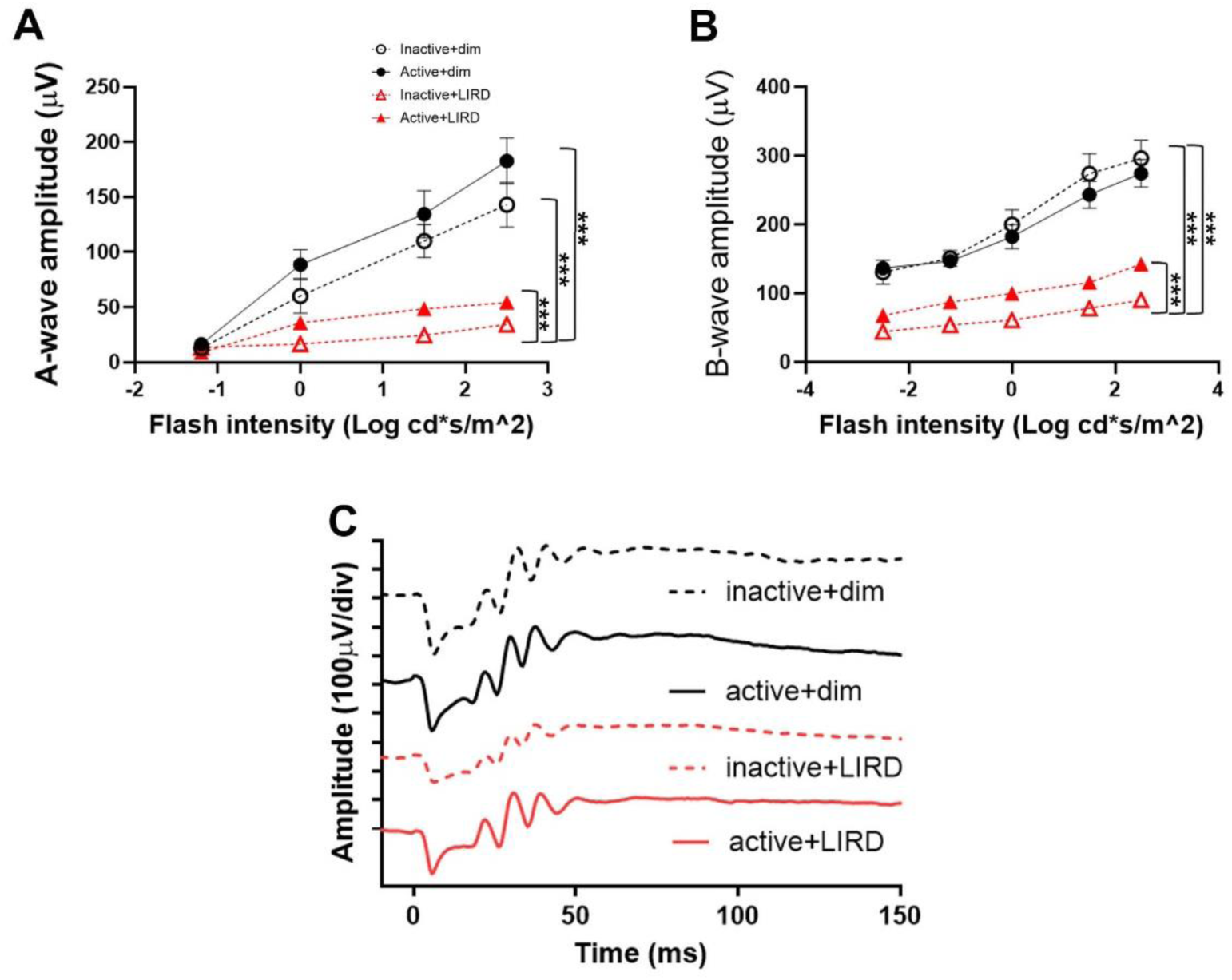
Active+LIRD mice have conserved retinal function 5 days post-LIRD. To confirm treadmill exercise preserved retinal function as demonstrated in our laboratories previously, electroretinography (ERG, **A,B**) recordings, quantifying a- and b-wave amplitudes were performed to noninvasively measure retinal function. Representative ERG waveforms shown were collected from maximum dark-adapted stimuli (1.5 log cd s/m^2^,**C**). Active+LIRD mice had significant preservation of a-wave (A) and b-wave (B) amplitudes compared to inactive+LIRD mice. A- and b-waves show rod photoreceptor and inner retina function, respectively. Two-way ANOVA with Tukey’s multiple comparison analysis was performed. N=10 per group, **p<0.001, values are mean±SEM.

### Treadmill exercise preserves photoreceptor nuclei 1- and 5-days post-LIRD and reduces photoreceptor apoptosis

To observe the effects of treadmill exercise in preserving photoreceptor nuclei density, morphometric quantifications were performed on hematoxylin and eosin (H&E) stained retinal sagittal sections. Initially following LIRD (1-day post-LIRD, **Figure 2A, B**), there was a significant decrease in the number of photoreceptor nuclei in the ONL in LIRD groups compared to dim groups (p<0.001). By 5 days post-LIRD, inactive+LIRD animals had a significant reduction in photoreceptor density compared to active+LIRD animals (**Figure 2C, D**; p<0.001). TUNEL staining revealed 1-day post-LIRD inactive+LIRD animals had a significant increase of photoreceptor apoptosis compared to active+LIRD (**Figure 3A, B**), with this trend continuing to 5-days post-LIRD (**Figure 3C, D**).

**Figure 2.**
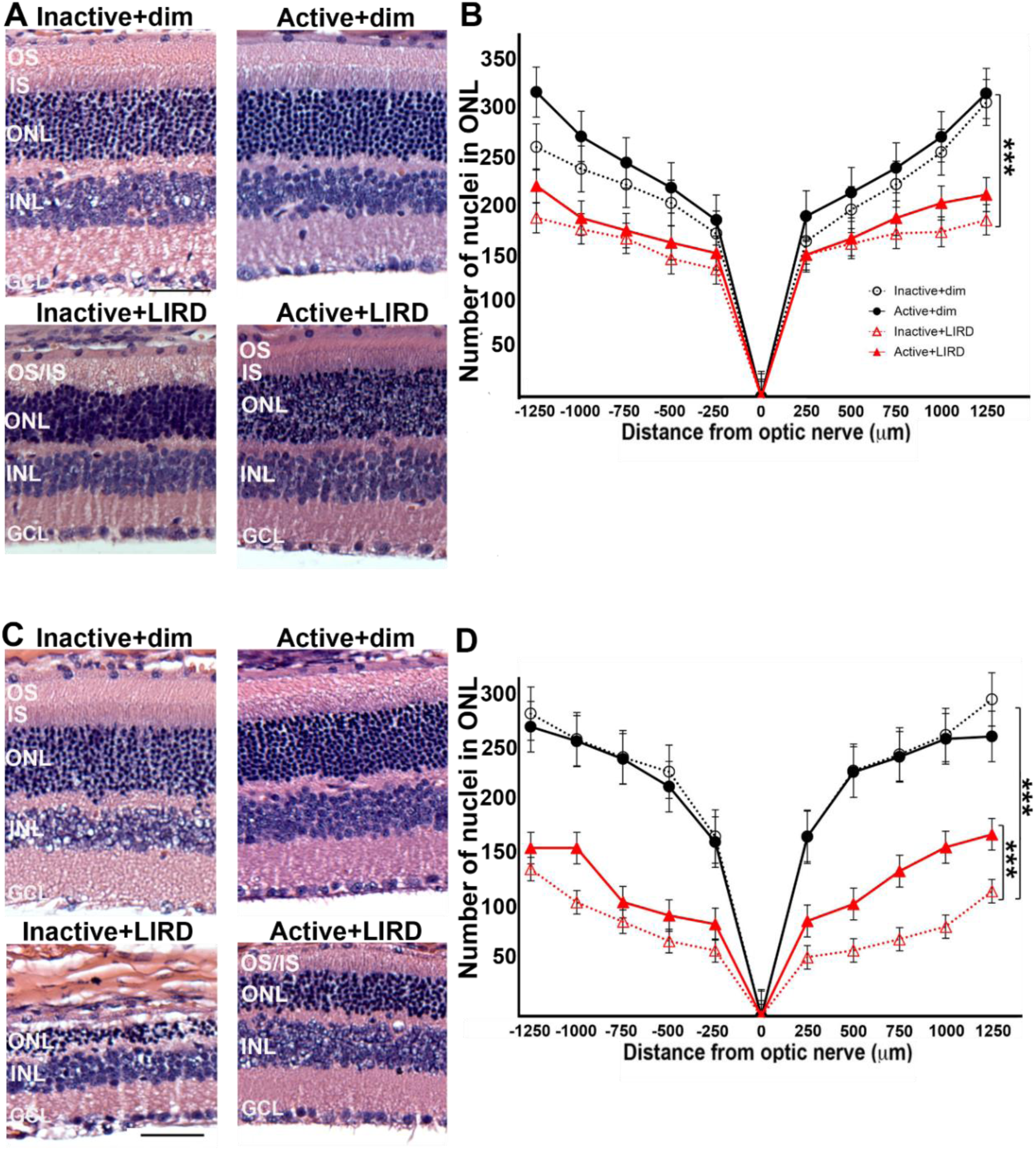
Treadmill exercise preserves photoreceptor nuclei 1 and 5 days post-LIRD. Retinal sections from all experimental groups (**A, C**) were used to quantify photoreceptor nuclei present in the outer nuclear layer (ONL). Morphometric analyses were constructed by plotting the quantification of the ONL nuclei at 1 and 5 days post-LIRD (**B, D**). 1 day post LIRD, photoreceptor nuclei numbers start to decline in both LIRD groups compared to dim groups and after 5 days, there is a significant decrease in inactive+LIRD animals compared to active+LIRD animals (****p<0.0001). Retinal layers are as follows: outer segment (OS), inner segment (IS), outer nuclear layer (ONL), inner nuclear layer (INL) and ganglion cell layer (GCL). N=9 per group, 3 retinal sections quantified per animal, ****p<0.0001. Scale bar= 25μm, values are mean±SEM.

**Figure 3.**
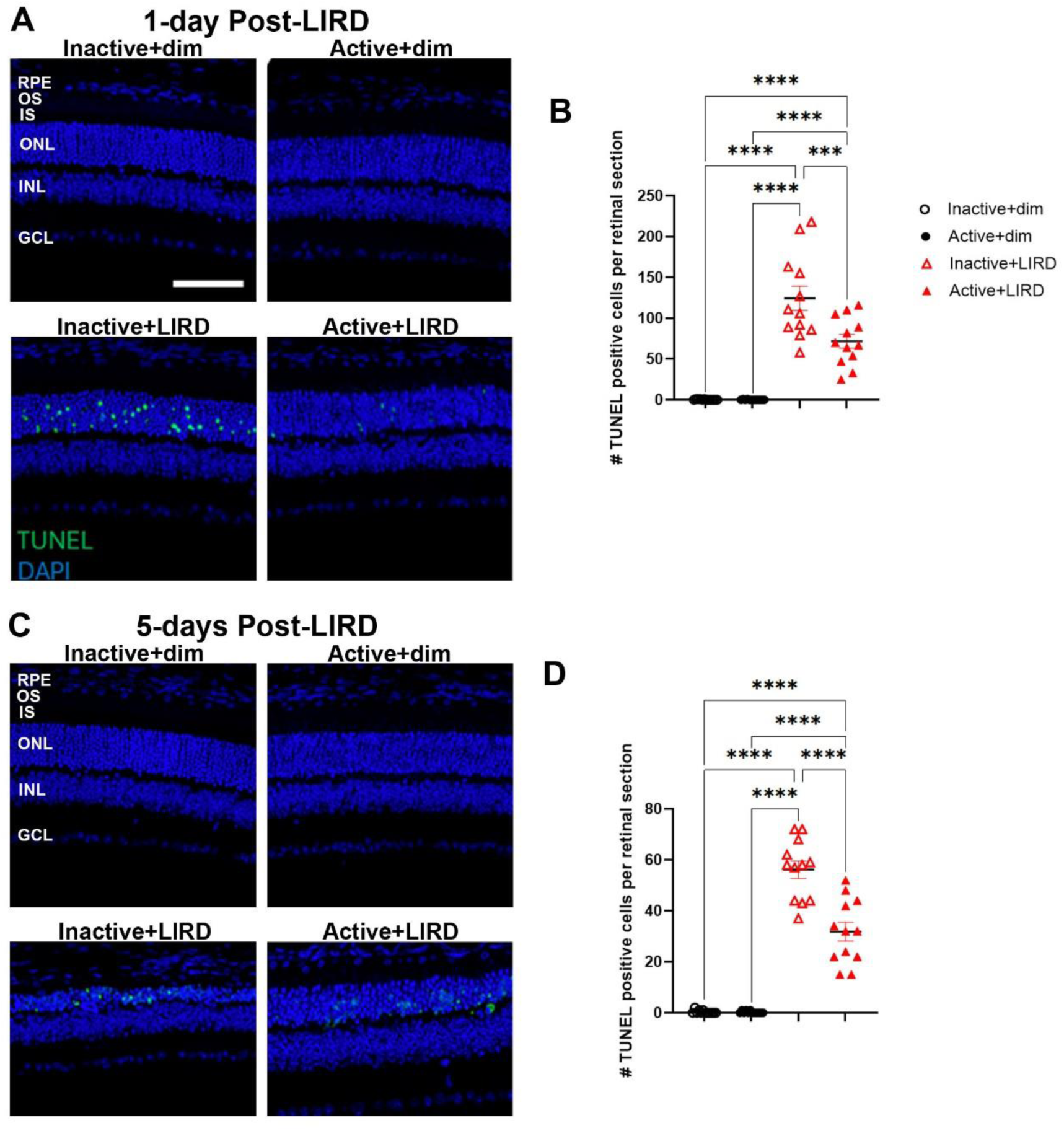
Active+LIRD mice have decreased photoreceptor apoptosis at 1- and 5 days post-LIRD. Quantification of TUNEL-positive cells (green) in the outer nuclear layer (ONL) revealed increased cell death occurred in inactive+LIRD animals compared to active+LIRD animals at 1- and 5-days post-LIRD. N=12 per group, 3 retinal sections quantified per animal, ***p<0.001, ****p<0.0001. Scale bar= 25μm, values are mean±SEM; blue, DAPI.

### Exercise partially suppressed LIRD-induced astrocyte and Müller glia activation

In order to quantify the reactivity of astrocytes and Müller glia, retinal sections from all experimental groups were labeled for glial fibrillary acidic protein (GFAP; **Figure 4A-D**) to positively label astrocytes and Müller glia as well as ionized calcium-binding adaptor molecule-1 (Iba1; **Figure 5A-D**) to positively label microglia. Retinal layer quantifications 1-day post-LIRD (**Figure 4A-D**) revealed there was no significant difference between active+LIRD and inactive+LIRD groups GFAP labeling, quantifying the inner nuclear layer (INL), inner plexiform layer (IPL) and ganglion cell layer (GCL). By 5days post-LIRD, there was a significant increase in GFAP labeling (**Figure 4A-D**) in the INL and IPL, although GCL was not statistically significant.

**Figure 4.**
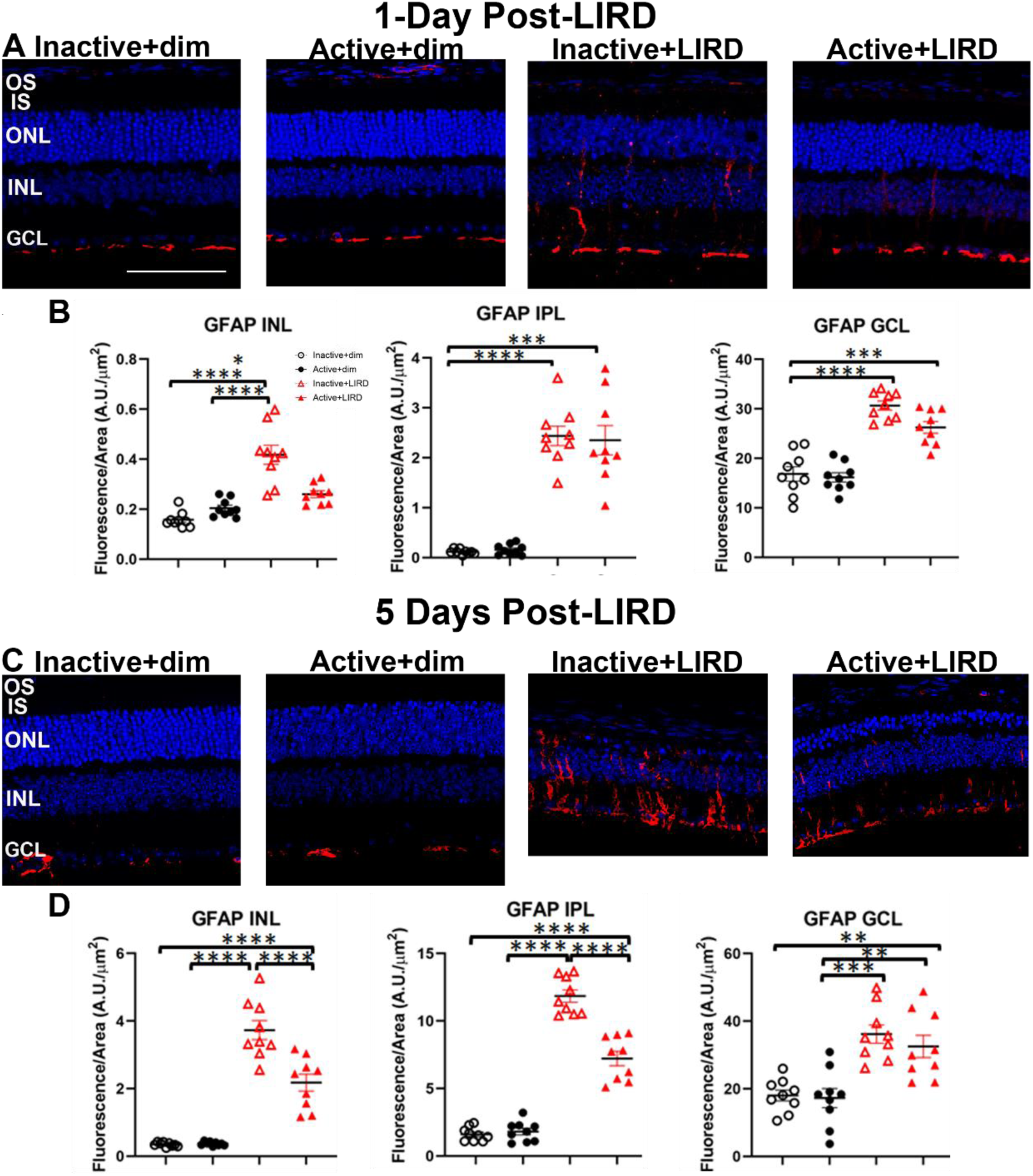
Exercise partially suppressed LIRD-induced astrocyte and Müller glia activation. Glial fibrillary acidic protein (GFAP, red) labeling in retinal sections from all experimental groups 1 and 5 days post-LIRD revealed active+LIRD mice had substantially less reactive astrocyte and Müller glia compared to inctive+LIRD mice. N=9 per group, 3 retinal sections quantified per animal, **p<0.01, ***p<0.001, ****p<0.0001. Scale bar= 50μm, values are mean±SEM; blue, DAPI.

**Figure 5.**
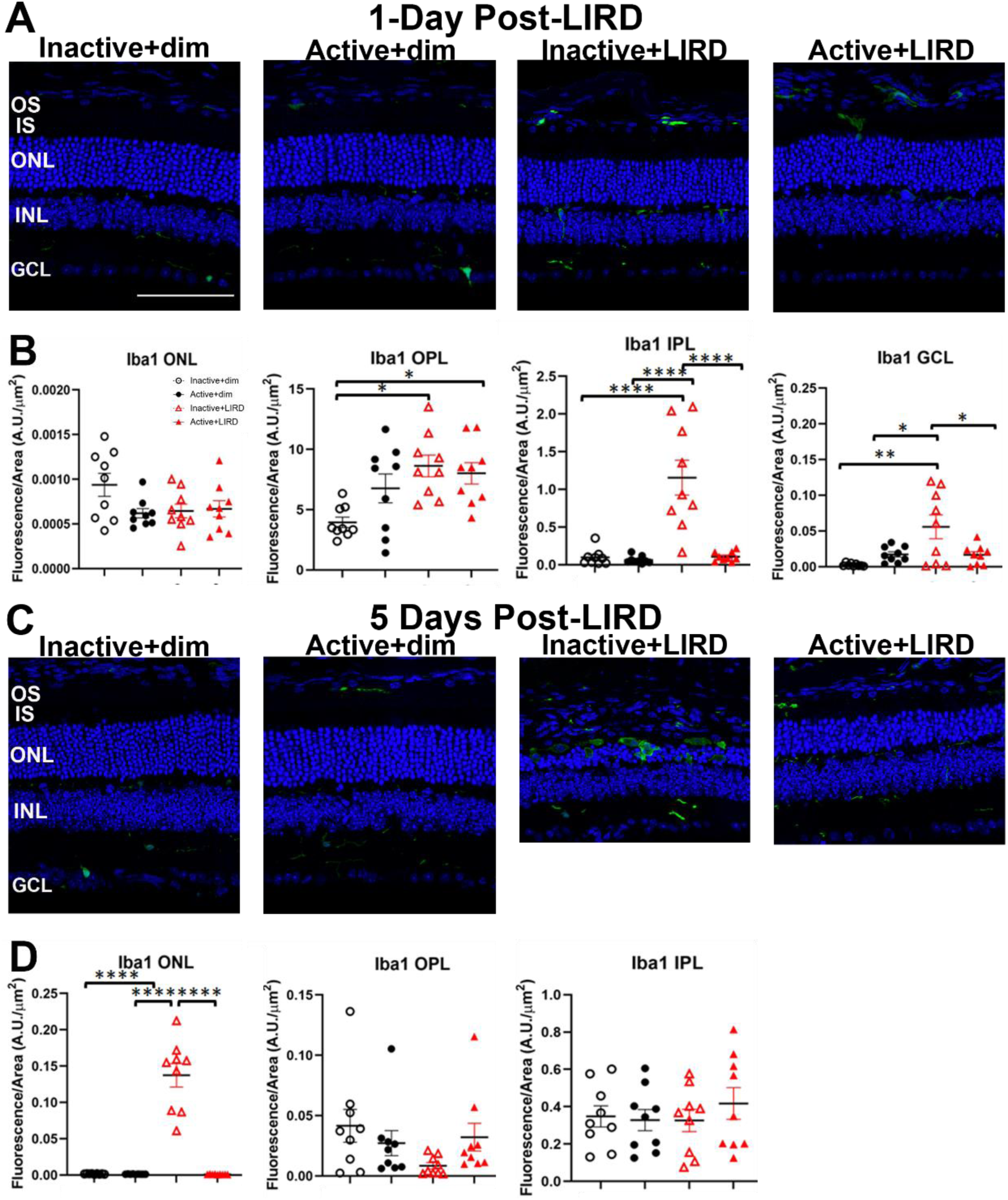
Active+LIRD mice exhibit fewer Iba1+ cells. Ionized calcium-binding adaptor molecule-1 (Iba1) immunofluorescence in retinal sections from experimental groups showed active+LIRD mice had decreased presence of microglia in the outer plexiform (OPL), inner plexiform (IPL) and ganglion cell layer (GCL) at 1day post-LIRD and the ONL and GCL at 5days post-LIRD compared to inactive+LIRD mice. N=9 per group, 3 retinal sections quantified per animal, *p<0.05, **p<0.01, ****p<0.0001. Scale bar= 50μm, values are mean±SEM; blue, DAPI.

### Active+LIRD mice have decreased presence of Iba1+ cells in neural retina

In order to quantify the presence of inflammatory glial cells, such as microglia, retinal sections from all experimental groups were labeled for ionized calcium-binding adaptor molecule-1 (Iba1; **Figure 5A-D**) which positively labels microglia and or macrophages. 1-day post-LIRD, there was increased Iba1 labeling in the IPL and GCL observed in inactive+LIRD animals compared to active+LIRD. In the outer plexiform layer (OPL), inactive+LIRD animals had an increased presence of Iba1+ cells compared to active+LIRD, although it was not significant (**Figure 5A, B**). By 5-days post-LIRD, inactive+LIRD animals had an increased presence of Iba1+ cells in the ONL and GCL compared to active+LIRD animals (**Figure 5C, D**).

### miRNA analyses reveal exercise promotes cell survival, oxidative stress regulation and photoreceptor homeostasis

To quantify gene expression levels between treatments (exercised vs. control and dim vs. LIRD), a MicroArray assay was conducted on retinal miRNA 1-day and 5-days post-LIRD, with significant changes in expression plotted on volcano graphs **(Figure 6A-D)**. Between active+dim and inactive+dim groups 1-day post-LIRD, miR-17 and miR-320 were downregulated, while miR-1224 and miR-183-5p were upregulated **(Figure 6A)**. In comparison, 1-day post-LIRD active+LIRD and inactive+LIRD groups showed downregulation of miR-135a-1-3p and miR-451b, upregulation of miR-1199 and miR-192-5p **(Figure 6B)**. miRNA expression 5-days after induced retinal damage was also analyzed for more long-term effects. Active+dim and inactive+dim groups 5-days post-LIRD showed downregulation of miR-3065 and upregulation of miR-1298-5p, miR-99b-5p, and miR-139 **(Figure 6C)**. Compared to the active+LIRD and inactive+LIRD groups 5-days post-LIRD, miR-302b-5p and miR-187 were significantly upregulated **(Figure 6D)**.

**Figure 6.**
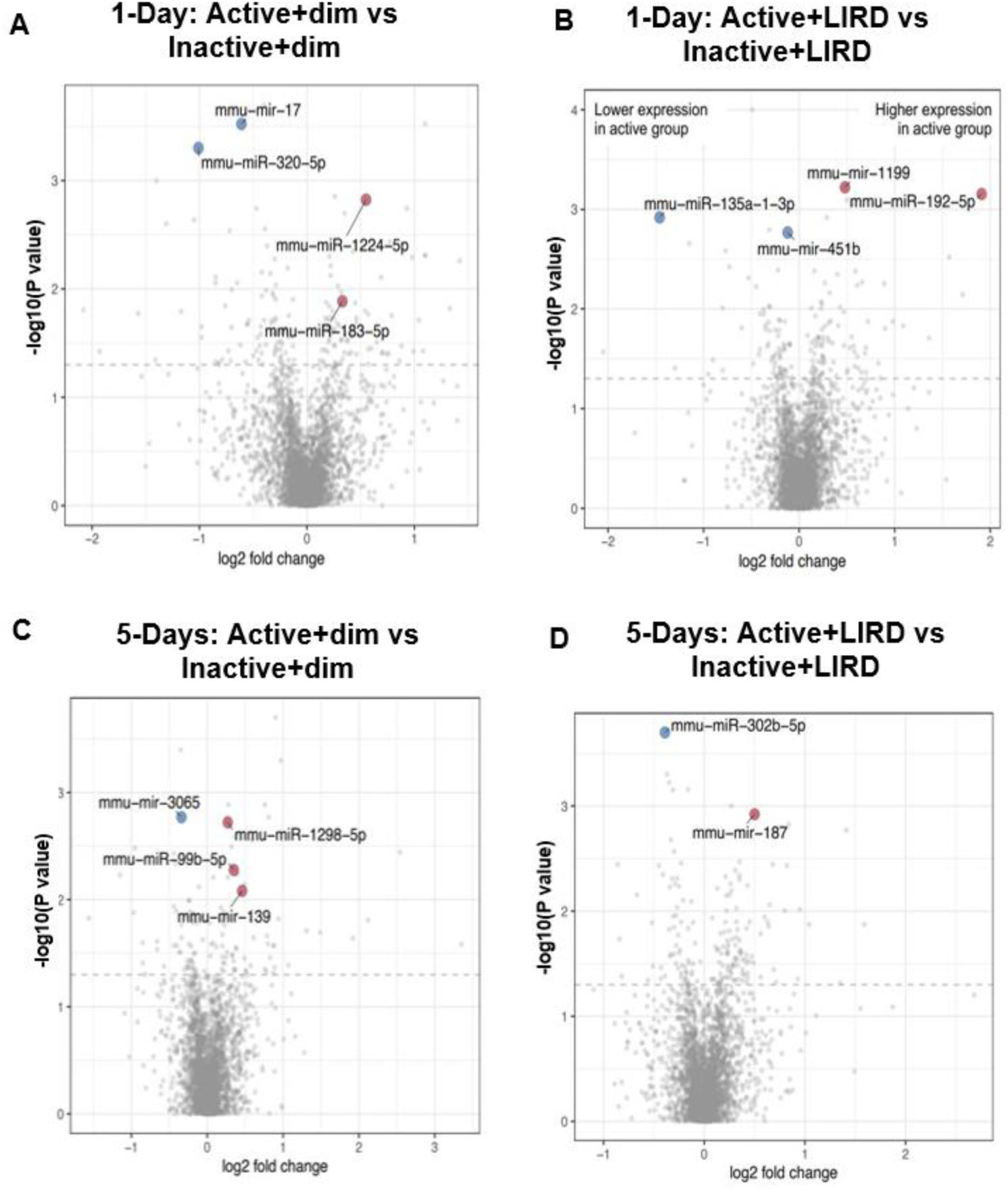
miRNA analyses reveal exercise promotes cell survival, oxidative stress regulation and photoreceptor homeostasis. Microarray analysis revealed exercise alters miRNA expression comparing dim and LIRD groups at 1- and 5-days. Altered miRNA expression profiles in active groups are associated with cell survival (miR-302b), oxidative stress regulation (miR-192-5p) and photoreceptor homeostasis (miR-187).

## Discussion

Exercise as a neuroprotective intervention has long been supported as treatment for neurodegenerative conditions like Alzheimer’s and Huntington’s disease, as it helps protect against DNA damage, protein misfolding and dysfunction, and apoptosis^12^. Studies from our group and others have shown exercise helps preserve photoreceptor function, structure, vasculature, astrocyte plasticity and inflammatory response^3,13,14,4,5^. Additionally, previous research in Alzheimer’s disease mouse models have shown that regular exercise reduces ocular NF-κB-induced inflammation and upregulates molecular markers (FNDC5, PGC-1α, and SIRT1) involved in mitochondria biogenesis and cell repair in the retina^15^. Similarly, aerobic exercise-induced brain derived neurotrophic factor (BDNF) expression has also been shown to provide neuroprotective measures by stabilizing photoreceptor nuclei, promoting astrocyte plasticity and in axotomized optic nerve mouse models, salvaging damaged retinal ganglion cells^13,3,16^. However, most notably, physical activity protects against neuronal apoptosis. For example, by suppressing pro-apoptotic proteins and enhancing the expression of cell survival factors through treadmill or running wheel exercise, an anti-apoptotic effect can be observed in rodent retinal cells^5,17^. Our study aims to add to these findings by investigating how exercise modulates cell death, inflammatory response, and miRNA expression and the inflammatory response for retinal neuroprotection 1-day and 5-days post LIRD.

This study finds that regular exercise promotes neuroprotective effects in mice with LIRD. After 2 weeks of 60-minute daily treadmill exercise, active+LIRD mice showed preserved photoreceptor density, decreased ONL cell apoptosis, suppression of inflammatory markers in retinal glial cells, and altered of miRNAs (miR-302b, miR-192-5p, miR-187) related to inflammatory responses, compared to inactive+LIRD mice 1-day and 5-days after LIRD. Our results provide support for exercise-based therapies to treat and provide protective, preventative measures against neurodegeneration and neuroinflammatory diseases.

Similar to our previous studies, electroretinography and TUNEL staining analyses revealed exercised mice had preservation of retinal cell function and a reduction in photoreceptor apoptosis compared to inactive+LIRD mice. We found 5-days post-LIRD, active+LIRD mice consistently showed significantly higher a- and a-wave amplitudes on electroretinogram graphs. Higher a- and b-wave amplitudes, a marker of the strength of rod photoreceptor cells’ electrical signals, indicates exercise has a preservative effect on photoreceptor cell signaling in active+LIRD mice, even after induced damage. In contrast, a diminished a- and b-wave photoreceptor response in inactive+LIRD mice indicates a weakened photoreceptor response, and is consistent with our previously reported results, which are comparable to those with retinal degenerative disorders such as diabetic retinopathy^3,4,11,13,18^. Our experiments show that active+LIRD mice maintain functional a-wave amplitudes 1.98x the amplitudes of inactive+LIRD mice, and b-wave amplitudes 1.72x the amplitudes of the inactive+LIRD mice, suggesting preserved retinal function even after light-induced degeneration due to treadmill exercise.

Additionally, H&E-stained sagittal sections and TUNEL analyses conducted 1-day and 5-days post-LIRD showed preserved photoreceptor nuclei in the outer nuclear layer (ONL) in active+LIRD mice compared to inactive+LIRD mice. H&E stains after LIRD shows that the mice who were treadmill-active had significantly less ONL nuclear degeneration throughout the retina than those who were not treadmill-active. This is supported by the fact that there was a significantly greater amount of apoptotic cells detected by TUNEL in the inactive+LIRD mice as opposed to the active+LIRD groups. It is important to note that although they still experienced nucleic degeneration from toxic light, active+LIRD mice still retained a significantly higher surviving cell density than the inactive+LIRD group. With a higher density of surviving photoreceptors and decreased quantity of photoreceptor apoptosis 1-day and 5-days post-LIRD exposure, active mice have noticeably healthier, preserved retinal layers in contrast to inactive mice.

Neuroprotective effects like those described above may be triggered by exercise’s anti-inflammatory benefits. It has been determined that retinal inflammation drives neurodegeneration, especially through the upregulation of inflammatory cytokine and chemokine pathways such as NF-κB and members of the IL-1 family of pro-inflammatory interleukins by activated retinal microglia, macrophages, astrocytes, and Muller glial cells^19,20^. Accumulation of reactive microglia and astrocytes is characteristic of retinal inflammation-induced retinal pathologies such as age-related macular degeneration (AMD), as well as typical aging^3,21^. However, previous research indicates that exercise can mitigate the inflammatory response within the retina by reducing levels of pro-inflammatory cytokine and chemokine expression (KC and IP-10) involved with retinal vasculature as well as mitigating astrocyte morphology and gene expression^3,4^. In this study, we further explore the role of inflammatory markers through retinal glial cell recruitment and inflammation.

Although they are mainly known for providing structural and cellular support, astrocytes and Muller glia (collectively called macroglia) also play a role in the inflammatory response through the release of cytokines and chemokines throughout the retinal layer^22–24^. In mammalian cells, upregulation of chemokine release by Muller glia is correlated with retinal stress, damage, and disease, especially in inflammatory diseases like proliferative vitreoretinopathy^25,26^ . However, they are also known to promote neuroprotection through the release of growth factors such as BDNF and nerve growth factor (NGF), and Muller glia derived progenitor cells are still currently being studied for their neuroregenerative potential^27^. For example, zebrafish and chick retinas have the potential to regenerate retinal neurons and glia by inducing stem cell-like properties in response to damage^28–30^. In a similar supportive role, astrocytes are heavily involved in regulating inflammatory cascades and supporting survival and axonal regeneration after induced damage^31,32^. Because of their collective roles in the immune response, macroglia are often used as markers to quantify retinal condition after inflammation or damage.

In our study, we found that exercise partially suppresses the recruitment of pro-inflammatory, reactive macroglia 1-day and 5-days after light-induced damage (fig. 4). Active+LIRD mice showed a significant reduction in astrocyte and Muller glia labeling of astrocytes and Muller glia in the INL and IPL compared to inactive+LIRD mice, indicating that exercise may mitigate the activation of pro-inflammatory glia. In addition, Iba1+, a microglia and macrophage marker, was also less present in the ONL, IPL, and GCL in active+LIRD mice versus inactive+LIRD. This decreased inflammatory glial cell response in active+LIRD mice indicates that exercise plays a role in suppressing the detrimental inflammatory response, allowing for increased protection against inflammation-induced neurodegeneration.

MicroRNAs, or miRNAs, are also an important factor in regulating cells’ inflammatory response^33^. miRNAs are non-coding RNAs that primarily control gene expression in response to environmental influences and even help time retinal neurogenesis^34,35^. In canine retinitis pigmentosa models, anti-apoptotic miRNAs have been found to be upregulated and pro-apoptotic miRNAs downregulated to potentially counteract retinal degeneration^36^. In Muller glial cells, inflammation associated with diabetic retinopathy leads to the upregulation of certain immune-related genes, and other types of damage contribute to promoting oxidative stress and retinal inflammatory factor expression^37^. Our study further confirms the conclusion that miRNAs are a significant factor in determining retinal cell survival. We found that 1-day and 5-day post-LIRD combined with treadmill exercise significantly affects the expression of miR-192-5p and miR-187, which are involved in apoptosis and the retinal inflammatory response, while downregulating the expression of miR-302b, a miRNA involved in the progression of aging and inflammation. Even in control mice without light-induced retinal degeneration, changes in expression after exercise were observed in miRNAs involved in regulating angiogenesis and cell proliferation, both 1-day and 5-days post-LIRD.

In groups not treated with LIRD, mice were kept in a dim environment and randomly sorted into active or inactive groups for treadmill exercise. Between active+dim and inactive+dim groups 1-day post-LIRD, upregulation of miR-1224 was observed, a miRNA involved in suppressing glioma^38^ and improving post-stroke neuronal vasculature^39^. miR-183-5p, a miRNA modulating retinal function, was also upregulated^40^. In contrast, miR-17 and miR-320, anti-angiogenesis and anti-survival factors were downregulated^41–44^. We found similar results 5-days post-LIRD. Tumor-suppressing and apoptotic genes^45–47^, miR-1298-5p and miR139 were upregulated in addition to retinal functionality gene miR-99b-5p^48^ while miR-3065, a factor involved in stemness and colorectal cancer metastasis, was downregulated^49^. Taken together, it can be suggested that even without being affected with retinal degeneration, treadmill exercise provides neuroprotective effects by upregulating miRNAs involved in angiogenesis and tumor suppression, and downregulating miRNAs involved in cancer proliferation.

The function of miR-192-5p has been previously explored in diseases like cancer and digestive disorders, and play vastly different roles based on the affected system^50^.One study reported a significant reduction in miR-192-5p expression in retinal microvascular endothelial cells from diabetic retinopathy (DR) patients^51^.It was found that overexpression of miR-192-5p helped suppress cognitive dysfunction in mice with depression and was correlated with greater endothelial cell proliferation and angiogenesis in DR patients^50^. Studies conducted on other neurodegenerative diseases showed similar results, in which miR-192-5p expression was downregulated in patients with AMD^52^. Upregulation of miR-192-5p has also been found to play a role in inhibiting metastatic potential of colon cancer by downregulating expression of Bcl-2, Zeb2 and VEGFA, factors that inhibit apoptosis and promote cell growth. Based on these cumulative results, it can be suggested that the miR-192-5p acts a as a modulator of vascular endothelial growth factor^51,53^. In this study, we found there was significant upregulation of miR-192-5p in the active+LIRD group compared to the inactive+LIRD group. Essentially, in mouse models with light-induced retinal degeneration, forced treadmill exercise is correlated with upregulation of miRNAs involved in vascular endothelial growth factors in the retina, potentially counteracting neural degeneration through supportive angiogenesis^14^.

miR-187 is a microRNA also found to contribute to cell survival in the retina. Decreased expression of miR-187 has been reported to correlate with increased expression of Smad7, a protein involved in inducing apoptosis in retinal ganglion cells. Similarly, a deficit of miR-187 leads to oxidative stress-induced cellular apoptosis, and thus neurodegeneration ^54,55^. In this experiment, we found that 1-day and 5-day active+LIRD mice displayed upregulation of miR-187 post-exercise routine, compared to inactive+LIRD mice. This increased miR-187 expression could contribute to neuroprotection by resisting apoptotic signals triggered by oxidative stress, suggesting treadmill exercise as a viable treatment for preventing excess neuronal death/degeneration.

Additionally, it is also important to note the downregulation of expression of miR-302b-5p in active+LIRD mice compared to the inactive+LIRD mice. miR-302b is part of the miR-302/367 cluster, a family of microRNAs involved in cell survival and pluripotency. Recently, this cluster has been highly studied for its role in cancer treatment, and it has been found that regulating the expression of this cluster can suppress the development of tumors and control growth/apoptosis signals^56^. Interestingly, miR-302b-5p seems to play a role in both tumor suppression and promoting proliferation. In a study on esophageal squamous cell carcinoma, miR-302b was shown to downregulate expression of ErbB4, a tumor growth and metastatic factor, as well as induce apoptosis in cancer cells^57^. In another study on human adipose tissue-derived mesenchymal stem cells (hADSCs), miR-302b was analyzed under the context of protecting hADSCs from oxidative stress-induced apoptosis and promoting cell proliferation by downregulating *CDKN1A*/p21 (a cell cycle disruptor)^58^. Taking into account the various regulatory functions of miR-302b, its role in retinal degeneration can be inferred. Because the downregulation of miR-302b-5p in active+LIRD mice correlates with increased cell survival in the context of this experiment, this data suggests that miR-302b-5p has a potential role in inhibiting apoptosis. By downregulating a factor that induces apoptosis in response to stress, cell survival is promoted, as we report in the active+LIRD animals through increased ONL counts and a decreased presence of TUNEL positive cells.

In this study, we demonstrate that treadmill exercise has a neuroprotective effect against retinal degeneration. Compared to inactive+LIRD mice at 1- and 5-day checkpoints, active+LIRD mice showed a significantly preserved density of retinal photoreceptors, slowed apoptosis in the ONL, decreased recruitment of inflammatory cells in the retinal layer, and modified expression of miR-192-5p, miR-187, and miR-302b to promote cell survival and growth. Even in mice not treated with LIRD, upregulation of neuroprotective miRNAs (miR-1224-5p, miR-183-5p, miR-1298-5p, miR139, miR-99b-5p) and downregulation of cancer-related miRNAs (miR-17, miR-320-5p, miR-3065) was observed between active+dim and inactive+dim groups. Our results align with published literature claiming that consistent lifelong physical activity is associated with better quality of late-life vision, as well as a later onset of vision-related disease, such as retinitis pigmentosa, glaucoma, and AMD ^59–61^. In addition to supporting previous scientific research, our study also finds novel connections between specific regulatory miRNAs and neuroprotection. This work highlights how physical exercise can either up- or down-regulate specific miRNAs that promote retinal neuroprotection. However, as the experiment was more short-term focused, the long-term effects of exercise on immunoregulation and miRNA modulation on the retina may need to be explored, as well asdifferences in regulation across retinal degenerative diseases. Future directions may include trying to find methods of exogenous incorporation of miRNA into the body to promote neuroprotection in lieu of exercise, or creating optimized strategies for physical activity-related therapies. Overall, this study finds evidence for the neuroprotective benefits of exercise, and lends support for exercise-based therapies to reduce retinal neurodegeneration, inflammation, and importantly, highlights differences in miRNA expression initially after retinal insult as well as 5-days post retinal damage.

## Availability of data and materials

miRNA data are available on GEO, accession number XXX. Although these data are not currently publicly available for sharing, requests for sharing can be sent to the Corresponding Author and will be evaluated on an individual basis.

## List of abbreviations

| Abbreviation | Expanded |
| --- | --- |
| AMD | Age-related macular degeneration |
| ANOVA | Analysis of Variance |
| ARVO | Association for Research in Vision and Ophthalmology |
| BALB/c | Bagg Albino /c strain |
| Bcl-2 | B-cell lymphoma 2 |
| BDNF | Brain derived neurotrophic factor |
| CDKN1A/p21 | Cyclin-dependent kinase inhibitor 1A |
| CEL | Cell intensity file |
| DAPI | 4',6-diamidino-2-phenylindole |
| DNA | Deoxyribonucleic acid |
| DR | Diabetic retinopathy |
| ErbB4 | Receptor tyrosine-protein kinase ErbB-4 |
| ERG | Electroretinography |
| FNDC5 | Fibronectin type III domain-containing protein 5 |
| GCL | Ganglion cell layer |
| GFAP | Glial fibrillary acidic protein |
| H & E | Hematoxylin and eosin |
| hADSC | Human adipose tissue-derived mesenchymal stem cell |
| Iba1 | Ionized calcium-binding adaptor molecule 1 |
| IL-1 | Interleukin-1 |
| INL | Inner nuclear layer |
| IP | Intraperitoneal |
| IP-10 | Interferon gamma inducible protein-10 |
| IPL | Inner plexiform layer |
| IS | Inner segment |
| KC | Keratinocyte-derived chemokine |
| LED | Light emitting diode |
| LIRD | Light-induced retinal degeneration model |
| miR- | Micro RNA |
| miRNA | Micro RNA |
| NF-κB | Nuclear factor-kappa B |
| NGF | Nerve growth factor |
| ONL | Outer nuclear layer |
| OPL | Outer plexiform layer |
| OS | Outer segment |
| PBS | Phosphate-buffered saline |
| PGC-1α | Peroxisome proliferator-activated receptor-gamma coactivator 1-alpha |
| RD | Retinal degeneration |
| RMA | Robust multiarray average |
| RNA | Ribonucleic acid |
| ROUT | Robust regression and outlier removal |
| RPE | Retinal pigmented epithelium |
| SEM | Standard error of the mean |
| SIRT1 | Sirtuin 1 |
| Smad7 | Mothers against decapentaplegic homolog 7 |
| TAC | Transcriptome analysis console |
| TUNEL | Terminal deoxynucleotidyl transferase dUTP nick end labeling |
| VEGF | Vascular endothelial growth factor |
| Zeb2 | Zinc finger E-box-binding homeobox 2 |
| ZT3-5 | Zeitgeber Time 3-5 |

## Declarations

## Acknowledgements

Not applicable.

## Contributions

HBH, VC, TP, TEA, GSR, JCT, RN, AJF, JMN, MTP, JHB, KLB contributed substantially to study conception and design, data collection, analysis, and interpretation. HBH, TP, TEA,GSR, JCT, RN, AJF, JMN, MTP, JHB, KLB contributed to drafting the article or revising it critically for important intellectual content. All authors gave final approval for submission and publication.

## Ethics approval

All animal procedures were approved by the Atlanta VA Institutional Animal Care and Use Committee and conform to the Association for Research in Vision and Ophthalmology (ARVO) Statement for the Use of Animals in Ophthalmic and Vision Research.

## Consent for publication

Not applicable.

## References

1. Allen RS, Hanif AM, Gogniat MA, et al. TrkB signalling pathway mediates the protective effects of exercise in the diabetic rat retina. Eur J Neurosci. 2018;47(10):1254–1265. doi:10.1111/ejn.13909

2. Chu-Tan JA, Kirkby M, Natoli R. Running to save sight: The effects of exercise on retinal health and function. Clin Exp Ophthalmol. 2022;50(1):74–90. doi:10.1111/ceo.14023

3. Bales KL, Chacko AS, Nickerson JM, Boatright JH, Pardue MT. Treadmill exercise promotes retinal astrocyte plasticity and protects against retinal degeneration in a mouse model of light-induced retinal degeneration. J Neurosci Res. 2022;100(9):1695–1706. doi:10.1002/jnr.25063

4. Bales KL, Karesh AM, Hogan K, et al. Voluntary exercise preserves visual function and reduces inflammatory response in an adult mouse model of autosomal dominant retinitis pigmentosa. Sci Rep. 2024;14(1):6940. doi:10.1038/s41598-024-57027-9

5. Chu-Tan JA, Cioanca AV, Wooff Y, et al. Voluntary exercise modulates pathways associated with amelioration of retinal degenerative diseases. Front Physiol. 2023;14:1116898. doi:10.3389/fphys.2023.1116898

6. Zhang X, Girardot PE, Sellers JT, et al. Wheel running exercise protects against retinal degeneration in the I307N rhodopsin mouse model of inducible autosomal dominant retinitis pigmentosa. Mol Vis. 2019;25:462–476.

7. Kauppinen A, Paterno JJ, Blasiak J, Salminen A, Kaarniranta K. Inflammation and its role in age-related macular degeneration. Cell Mol Life Sci. 2016;73(9):1765–1786. doi:10.1007/s00018-016-2147-8

8. Proinflammatory Pathways Are Activated in the Human Q344X Rhodopsin Knock-In Mouse Model of Retinitis Pigmentosa - PubMed. Accessed January 28, 2025. https://pubmed.ncbi.nlm.nih.gov/34439829/

9. Das K, Rao LVM. The Role of microRNAs in Inflammation. Int J Mol Sci. 2022;23(24):15479. doi:10.3390/ijms232415479

10. Grimm C, Remé CE. Light damage as a model of retinal degeneration. Methods Mol Biol. 2013;935:87–97. doi:10.1007/978-1-62703-080-9_6

11. Mees LM, Coulter MM, Chrenek MA, et al. Low-Intensity Exercise in Mice Is Sufficient to Protect Retinal Function During Light-Induced Retinal Degeneration. Invest Ophthalmol Vis Sci. 2019;60(5):1328–1335. doi:10.1167/iovs.18-25883

12. Sujkowski A, Hong L, Wessells RJ, Todi SV. The protective role of exercise against age-related neurodegeneration. Ageing Res Rev. 2022;74:101543. doi:10.1016/j.arr.2021.101543

13. Hanif AM, Lawson EC, Prunty M, et al. Neuroprotective Effects of Voluntary Exercise in an Inherited Retinal Degeneration Mouse Model. Invest Ophthalmol Vis Sci. 2015;56(11):6839–6846. doi:10.1167/iovs.15-16792

14. Haupt HB, Nickerson JM, Boatright JH, Pardue MT, Bales KL. Effects of treadmill exercise on retinal vascular morphology, function, and circulating immune factors in a mouse model of retinal degeneration. Published online February 12, 2025:2025.02.11.637694. doi:10.1101/2025.02.11.637694

15. Okudan S, Sezer T, Tınkır Kayırbatmaz E, Belviranli M, Okudan N. Effect of regular exercise on ocular inflammation and mitochondrial biogenesis in experimental Alzheimer’s disease model. Cell Mol Biol (Noisy-le-grand). 2025;71(3):117–123. doi:10.14715/cmb/2025.71.3.14

16. He YY, Wang L, Zhang T, Weng SJ, Lu J, Zhong YM. Aerobic exercise delays retinal ganglion cell death after optic nerve injury. Exp Eye Res. 2020;200:108240. doi:10.1016/j.exer.2020.108240

17. Kim DY, Jung SY, Kim CJ, Sung YH, Kim JD. Treadmill exercise ameliorates apoptotic cell death in the retinas of diabetic rats. Molecular Medicine Reports. 2013;7(6):1745–1750. doi:10.3892/mmr.2013.1439

18. Kim MK, Aung MH, Mees L, et al. Dopamine Deficiency Mediates Early Rod-Driven Inner Retinal Dysfunction in Diabetic Mice. Invest Ophthalmol Vis Sci. 2018;59(1):572–581. doi:10.1167/iovs.17-22692

19. Palazzo I, Kelly L, Koenig L, Fischer AJ. Patterns of NFkB activation resulting from damage, reactive microglia, cytokines, and growth factors in the mouse retina. Exp Neurol. 2023;359:114233. doi:10.1016/j.expneurol.2022.114233

20. Nassar K, Grisanti S, Elfar E, Lüke J, Lüke M, Grisanti S. Serum cytokines as biomarkers for age-related macular degeneration. Graefes Arch Clin Exp Ophthalmol. 2015;253(5):699–704. doi:10.1007/s00417-014-2738-8

21. Vessey KA, Waugh M, Jobling AI, et al. Assessment of Retinal Function and Morphology in Aging Ccl2 Knockout Mice. Investigative Ophthalmology & Visual Science. 2015;56(2):1238–1252. doi:10.1167/iovs.14-15334

22. Drescher KM, Whittum-Hudson JA. Modulation of immune-associated surface markers and cytokine production by murine retinal glial cells. Journal of Neuroimmunology. 1996;64(1):71–81. doi:10.1016/0165-5728(95)00156-5

23. Bringmann A, Iandiev I, Pannicke T, et al. Cellular signaling and factors involved in Müller cell gliosis: Neuroprotective and detrimental effects. Progress in Retinal and Eye Research. 2009;28(6):423–451. doi:10.1016/j.preteyeres.2009.07.001

24. Yoo HS, Shanmugalingam U, Smith PD. Harnessing Astrocytes and Müller Glial Cells in the Retina for Survival and Regeneration of Retinal Ganglion Cells. Cells. 2021;10(6):1339. doi:10.3390/cells10061339

25. Norrie JL, Lupo M, Shirinifard A, et al. Latent Epigenetic Programs in Müller Glia Contribute to Stress, Injury, and Disease Response in the Retina. Published online October 17, 2023:2023.10.15.562396. doi:10.1101/2023.10.15.562396

26. Eastlake K, Banerjee PJ, Angbohang A, Charteris DG, Khaw PT, Limb GA. Müller glia as an important source of cytokines and inflammatory factors present in the gliotic retina during proliferative vitreoretinopathy. Glia. 2016;64(4):495–506. doi:10.1002/glia.22942

27. Fischer AJ, Reh TA. Müller glia are a potential source of neural regeneration in the postnatal chicken retina. Nat Neurosci. 2001;4(3):247–252. doi:10.1038/85090

28. Jadhav AP, Roesch K, Cepko CL. Development and neurogenic potential of Müller glial cells in the vertebrate retina. Progress in Retinal and Eye Research. 2009;28(4):249–262. doi:10.1016/j.preteyeres.2009.05.002

29. Simón MV, De Genaro P, Abrahan CE, de los Santos B, Rotstein NP, Politi LE. Müller glial cells induce stem cell properties in retinal progenitors in vitro and promote their further differentiation into photoreceptors. Journal of Neuroscience Research. 2012;90(2):407–421. doi:10.1002/jnr.22747

30. Fausett BV, Goldman D. A role for alpha1 tubulin-expressing Müller glia in regeneration of the injured zebrafish retina. J Neurosci. 2006;26(23):6303–6313. doi:10.1523/JNEUROSCI.0332-06.2006

31. Yoo HS, Shanmugalingam U, Smith PD. Harnessing Astrocytes and Müller Glial Cells in the Retina for Survival and Regeneration of Retinal Ganglion Cells. Cells. 2021;10(6):1339. doi:10.3390/cells10061339

32. White RE, Yin FQ, Jakeman LB. TGF-α increases astrocyte invasion and promotes axonal growth into the lesion following spinal cord injury in mice. Experimental Neurology. 2008;214(1):10–24. doi:10.1016/j.expneurol.2008.06.012

33. Saxena K, Rutar MV, Provis JM, Natoli RC. Identification of miRNAs in a Model of Retinal Degenerations. Invest Ophthalmol Vis Sci. 2015;56(3):1820–1829. doi:10.1167/iovs.14-15449

34. Zuzic M, Rojo Arias JE, Wohl SG, Busskamp V. Retinal miRNA Functions in Health and Disease. Genes (Basel). 2019;10(5):377. doi:10.3390/genes10050377

35. Conserved microRNA pathway regulates developmental timing of retinal neurogenesis | PNAS. Accessed June 30, 2025. https://www.pnas.org/doi/full/10.1073/pnas.1301837110

36. Genini S, Guziewicz KE, Beltran WA, Aguirre GD. Altered miRNA expression in canine retinas during normal development and in models of retinal degeneration. BMC Genomics. 2014;15(1):172. doi:10.1186/1471-2164-15-172

37. Jiang W, He S, Liu L, et al. New insights on the role of microRNAs in retinal Müller glial cell function. British Journal of Ophthalmology. 2024;108(3):329–335. doi:10.1136/bjo-2023-324132

38. Qian J, Li R, Wang YY, et al. MiR-1224-5p acts as a tumor suppressor by targeting CREB1 in malignant gliomas. Mol Cell Biochem. 2015;403(1):33–41. doi:10.1007/s11010-015-2334-1

39. Palakurti R, Biswas N, Roy S, et al. Inducible miR-1224 silences cerebrovascular Serpine1 and restores blood flow to the stroke-affected site of the brain. Mol Ther Nucleic Acids. 2023;31:276–292. doi:10.1016/j.omtn.2022.12.019

40. Zhang CJ, Xiang L, Chen XJ, et al. Ablation of Mature miR-183 Leads to Retinal Dysfunction in Mice. Invest Ophthalmol Vis Sci. 2020;61(3):12. doi:10.1167/iovs.61.3.12

41. Angiogenic microRNAs Linked to Incidence and Progression of Diabetic Retinopathy in Type 1 Diabetes | Diabetes | American Diabetes Association. Accessed July 16, 2025. https://diabetesjournals.org/diabetes/article/65/1/216/34936/Angiogenic-microRNAs-Linked-to-Incidence-and

42. Synchronous down-modulation of miR-17 family members is an early causative event in the retinal angiogenic switch | PNAS. Accessed July 16, 2025. https://www.pnas.org/doi/10.1073/pnas.1500008112

43. Tian B, Maidana DE, Dib B, et al. miR-17-3p Exacerbates Oxidative Damage in Human Retinal Pigment Epithelial Cells. PLoS One. 2016;11(8):e0160887. doi:10.1371/journal.pone.0160887

44. Sun N, Zhang D, Ni N, et al. miR-17 regulates the proliferation and differentiation of retinal progenitor cells by targeting CHMP1A. Biochemical and Biophysical Research Communications. 2020;523(2):493–499. doi:10.1016/j.bbrc.2019.11.108

45. Wang CM, Cheng BH, Xue QJ, Chen J, Bai B. MiR-1298 affects cell proliferation and apoptosis in C6 cells by targeting SET domain containing 7. Int J Immunopathol Pharmacol. 2017;30(3):264–271. doi:10.1177/0394632017720546

46. Stavast CJ, van Zuijen I, Erkeland SJ. MicroRNA-139, an Emerging Gate-Keeper in Various Types of Cancer. Cells. 2022;11(5):769. doi:10.3390/cells11050769

47. Stavast CJ, van Zuijen I, Karkoulia E, et al. The tumor suppressor MIR139 is silenced by POLR2M to promote AML oncogenesis. Leukemia. 2022;36(3):687–700. doi:10.1038/s41375-021-01461-5

48. Karali M, Persico M, Mutarelli M, et al. High-resolution analysis of the human retina miRNome reveals isomiR variations and novel microRNAs. Nucleic Acids Res. 2016;44(4):1525–1540. doi:10.1093/nar/gkw039

49. Li Y, Xun J, Wang B, et al. miR-3065-3p promotes stemness and metastasis by targeting CRLF1 in colorectal cancer. J Transl Med. 2021;19:429. doi:10.1186/s12967-021-03102-y

50. Frontiers | Emerging Role of MiR-192-5p in Human Diseases. Accessed June 30, 2025. https://www.frontiersin.org/journals/pharmacology/articles/10.3389/fphar.2021.614068/fu ll

51. fu XL, He FT, Li MH, Fu CY, Chen JZ. Upregulation of miR-192-5p inhibits the ELAVL1/PI3Kδ axis and attenuates microvascular endothelial cell proliferation, migration and angiogenesis in diabetic retinopathy. Published online July 26, 2022. doi:10.21203/rs.3.rs-1871717/v1

52. Ertekin S, Yıldırım Ö, Dinç E, Ayaz L, Fidancı ŞB, Tamer L. Evaluation of circulating miRNAs in wet age-related macular degeneration. Mol Vis. 2014;20:1057–1066.

53. Tang CZ, Yang JT, Liu QH, Wang YR, Wang WS. Up-regulated miR-192-5p expression rescues cognitive impairment and restores neural function in mice with depression via the Fbln2-mediated TGF-β1 signaling pathway. The FASEB Journal. 2019;33(1):606–618. doi:10.1096/fj.201800210RR

54. Zhang Q li, Wang W, Li J, Tian SY, Zhang TZ. Decreased miR-187 induces retinal ganglion cell apoptosis through upregulating SMAD7 in glaucoma. Biomedicine & Pharmacotherapy. 2015;75:19–25. doi:10.1016/j.biopha.2015.08.028

55. Zhang QL, Wang W, Alatantuya, et al. Down-regulated miR-187 promotes oxidative stress-induced retinal cell apoptosis through P2X7 receptor. International Journal of Biological Macromolecules. 2018;120:801–810. doi:10.1016/j.ijbiomac.2018.08.166

56. Liu J, Wang Y, Ji P, Jin X. Application of the microRNA-302/367 cluster in cancer therapy. Cancer Sci. 2020;111(4):1065–1075. doi:10.1111/cas.14317

57. Zhang M, Yang Q, Zhang L, et al. miR-302b is a potential molecular marker of esophageal squamous cell carcinoma and functions as a tumor suppressor by targeting ErbB4. Journal of Experimental & Clinical Cancer Research. 2014;33(1):10. doi:10.1186/1756-9966-33-10

58. Kim JY, Shin KK, Lee AL, et al. MicroRNA-302 induces proliferation and inhibits oxidant-induced cell death in human adipose tissue-derived mesenchymal stem cells. Cell Death Dis. 2014;5(8):e1385–e1385. doi:10.1038/cddis.2014.344

59. Levinson JD, Joseph E, Ward LA, et al. Physical Activity and Quality of Life in Retinitis Pigmentosa. Journal of Ophthalmology. 2017;2017(1):6950642. doi:10.1155/2017/6950642

60. Lee MJ, Wang J, Friedman DS, Boland MV, Moraes CGD, Ramulu PY. Greater Physical Activity Is Associated with Slower Visual Field Loss in Glaucoma. Ophthalmology. 2019;126(7):958–964. doi:10.1016/j.ophtha.2018.10.012

61. Past physical activity and age-related macular degeneration: the Melbourne Collaborative Cohort Study | British Journal of Ophthalmology. Accessed June 30, 2025. https://bjo.bmj.com/content/100/10/1353

